# Mixotrophy in phytoplankton: Prevalent use of organic compounds and lineage-specific variations suggest conserved ancestral traits

**DOI:** 10.64898/2026.02.04.703725

**Authors:** Nele Martens, Luisa Listmann, Jette Ludewigs, C.-Elisa Schaum

## Abstract

Mixotrophy is emerging as a default nutritional strategy in phytoplankton, but research seems so far isolated and mostly focusing on single phytoplankton groups or strains. Here we combined data from 24 oceanic and 22 freshwater strains to analyze phytoplankton’s ability to use dissolved organic compounds. The results emphasize that mixotrophy is ubiquitous in phytoplankton across functional groups and taxa isolated from various habitats and not strictly dependent on light or nutrient deficiencies. We propose that the prevalence of mixotrophy stems from phytoplankton’s heterotrophic ancestry, while the divergence into different lineages partly explains variation in mixotrophic behavior. Differences found in phytoplankton originating from different habitats may result from adaptive responses to habitat conditions, but conclusions need to be drawn carefully in line with the limited number of taxa included per habitat and potential niche partitioning. This study underscores the ecological significance of heterogeneous nutritional strategies and the necessity to address existing knowledge gaps to finally draw conclusions about the impact of mixotrophy in the context of single ecosystems and on global scales.

**Graphical abstract text:** Analyzing thousands of data points across 46 marine and freshwater strains, this study establishes mixotrophy as a ubiquitous metabolic strategy in phytoplankton. These findings challenge the autotrophic paradigm, demonstrating that organic matter acquisition is a default state with significant implications for our understanding of global carbon cycles.

**Graphical abstract:** 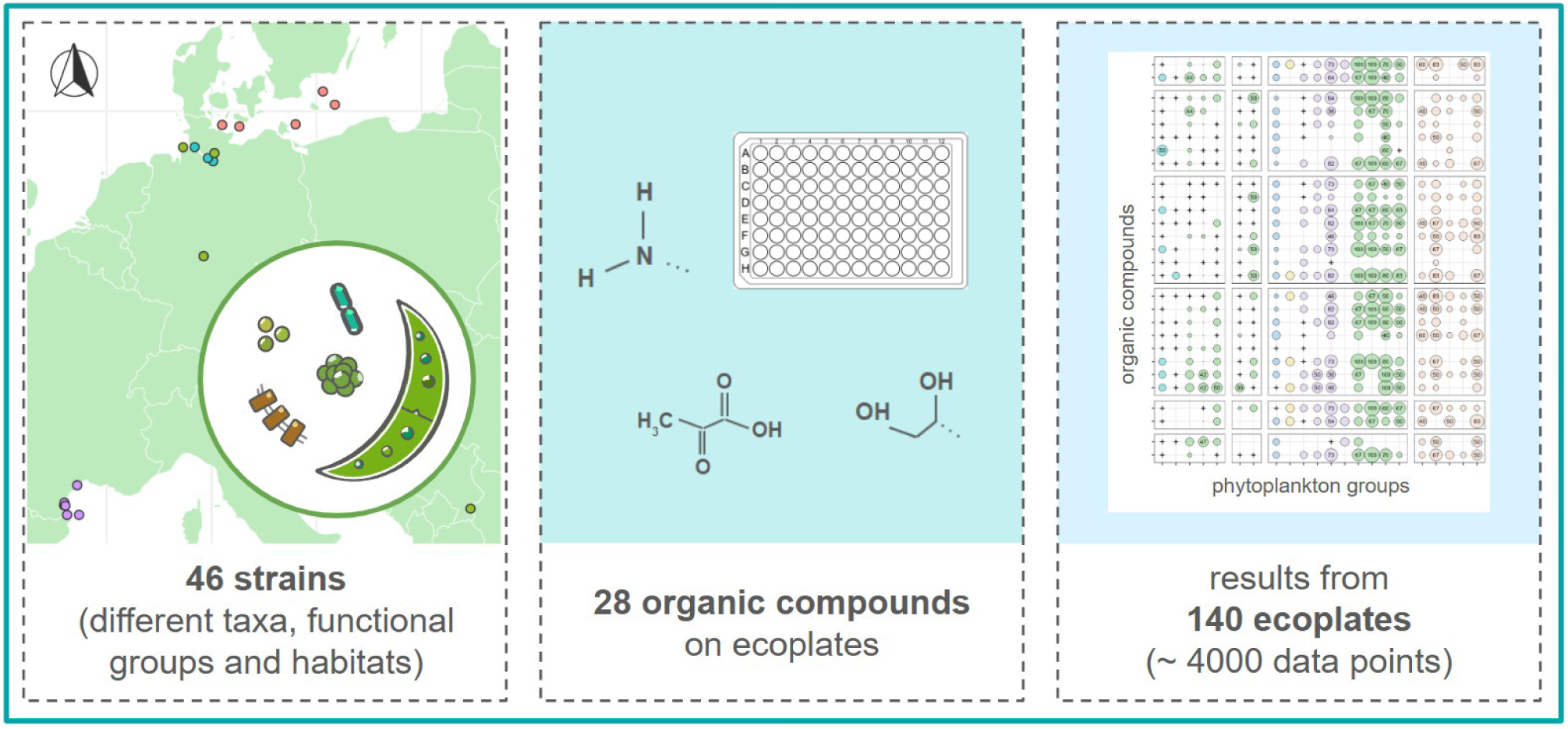

## 1. Introduction

Phytoplankton are still mostly considered autotrophs. However, recent studies challenge our understanding of the boundaries between different nutritional strategies among small aquatic organisms: Many phytoplankton taxa from various ecosystems have been shown to make use of organic material either by phagotrophy (Ahn & Glibert, 2024; Costa et al., 2025; Millette et al., 2023) or by using dissolved organic compounds such as sugars (Godrijan et al., 2022; Liu et al., 2024; Villanova & Spetea, 2021). The latter, also called osmotrophy, may be specifically widespread as it is not limited to certain morphotypes such as flagellates.

The use of organic matter affects the functioning of phytoplankton in ecosystems. For instance, exploiting heterotrophic pathways can help phytoplankton to survive and grow under nutrient- or light-limited conditions (Anderson et al., 2018; Calderini et al., 2022; Jones et al., 2009; Naselli-Flores & Barone, 2019; Tuchman et al., 2006) and hence drive photosynthesis and CO_2_ fixation. The use of organic matter by phytoplankton may also increase the transfer efficiency along food webs (Ward & Follows, 2016) and mixotrophic behavior in response to inorganic nutrient depletion can be linked to toxin production (Otte et al., 2025). How and the extent to which mixotrophic behavior affects communities and finally ecosystems and substrate cycles - as well as the potential impact on and of global change - remain widely unexplored. Exceptions are a few studies that largely deal with phagotrophy (Ahn & Glibert, 2024; Chen et al., 2024; Honig et al., 2025; Reinl et al., 2022; Wieczynski et al., 2023).

Phytoplankton use organic compounds with the aim of obtaining energy, harvesting N and P, or to directly use the compounds in other ways (e.g. to build proteins). As phytoplankton are able to synthesize required materials themselves, it has been suggested that mixotrophy is a strategy to deal with conditions impeding photosynthetic growth, such as light, nutrient, or even CO_2_ limitation (Calderini et al., 2022; Foresi et al., 2022; Li et al., 2021; Tuchman et al., 2006; Zhang et al., 2021). Yet, studies on phytoplankton’s mixotrophic behavior are often narrow, focusing on single compounds and strains covering single taxonomic groups and habitats.

Here we combined ecoplate data from 24 oceanic and 22 freshwater strains to analyze phytoplankton’s ability to use 28 different dissolved organic compounds. Specifically, we wanted to compare the results among taxa and ecosystems.

## 2. Methods

### Datasets and study areas

Our analysis comprises data from 140 ecoplates (see detailed description of laboratory procedures below) covering 46 phytoplankton strains (tab. S1) from different study areas. Those include the Baltic Sea, the Mediterranean Sea, the Elbe estuary, and different European bogs (fig. 1a, tab. S1).

**Fig. 1:**
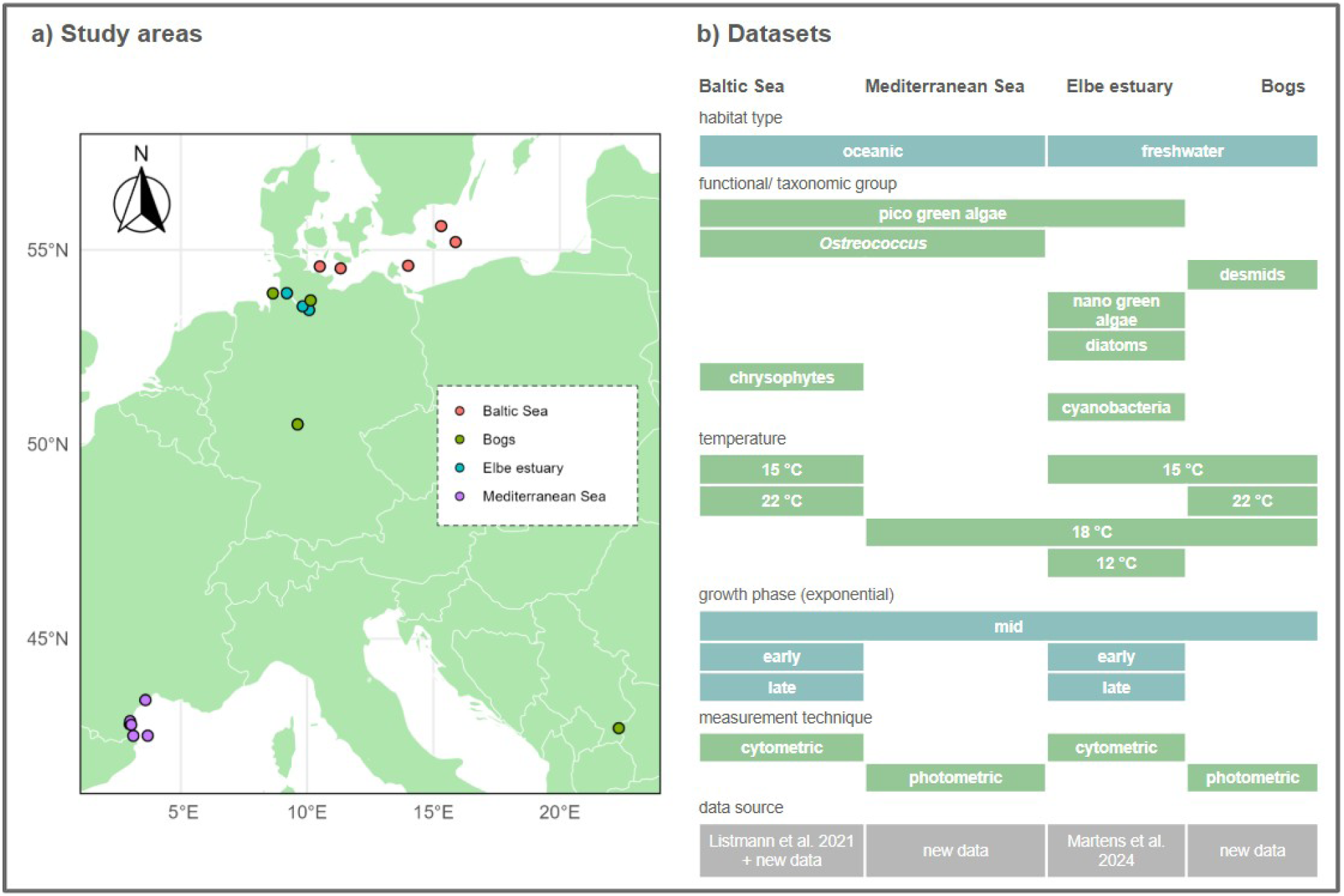
Overview of the included datasets. (a) shows the sampling stations from which the phytoplankton strains were isolated, (b) indicates where datasets differ and overlap with respect to additional parameters. Datasets are obtained from (Listmann et al., 2021; Martens et al., 2024) and unpublished data. See also table S1.

The Baltic Sea dataset is from previously published data (Listmann et al., 2021) and associated unpublished data. Chrysophytes and pico green algae (Chlorophyta < 3 µm) - including the key taxon *Ostreococcus* - were isolated from the Kiel area, Arkona Basin, and Bornholm Basin (fig. 1a, tab. S1). The Mediterranean Sea dataset contains new unpublished data. Strains of the pico green alga *Ostreococcus* were isolated from different lagoons, coastal areas and one open ocean station in the Gulf of Lion (fig. 1a, tab. S1). Strains were provided by the group of Gwenael Piganeau (Evolutionary and Environmental Genomics, Observatoire Océanologique de Banyuls-sur-Mer). The Elbe estuary dataset has been published in (Martens et al., 2024). Strains were mainly isolated from the area around Hamburg and a few from further downstream (fig. 1a, tab. S1). Strains include pico and nano green algae (Chlorophyta ∼ 3 - 20 µm), cyanobacteria, and a diatom (Bacillariophyta). The bog dataset is part of an unpublished study (Listmann et al., unpublished). It includes data of two desmids (Desmidiales; *Cosmarium botrytis* and *Closterium moniliferum*) - referred to as *Cosmarium* and *Closterium* throughout for clarity. Strains were provided by the Microalgae and Zygnematophyceae culture collection of the University of Hamburg (MZCH) (Von Schwartzenberg et al., 2013). They were isolated from different bogs in Germany and Serbia (fig. 1a, tab. S1). Note that throughout, for clarity, we consider the abovementioned groups (e.g. diatoms, desmids, pico green algae) functional groups (see also (Litchman et al., 2007)). They are, with the exception of pico and nano green algae, also distinct taxonomic groups, as indicated above.

The datasets partially differ in terms of experimental design and other laboratory-related factors. This is because the data were obtained with different objectives and not originally designed to be compiled. Figure 1b shows where datasets overlap with respect to certain aspects and where they differ (see also tab. S1). Each dataset contains a mostly unique set of taxa (tab. S1) and functional groups. Exceptions are the group of pico green algae (here Chlorophyta < 3 µm), which was included in the Baltic Sea, Mediterranean Sea and Elbe dataset, and, more particularly, *Ostreococcus*, which was included with multiple strains each in the Mediterranean Sea and Baltic Sea dataset (fig. 1b). Elbe and Baltic Sea strains were assessed at different time points during the exponential growth phase. Mediterranean Sea and bog strains were always assessed during the mid exponential phase (fig. 1b). Bog and Baltic Sea strains were assessed at different (treatment) temperatures (tab. S1). Mediterranean Sea and Elbe strains were always assessed at their incubation temperature (fig. 1b, tab. S1). Further, the measurement techniques differed for the datasets (fig. 1b, see also below and tab. S1).

### Laboratory procedures

We used Biolog ecoplates to analyze the potential use of organic compounds. Ecoplates are 96 well plates that contain different dissolved organic compounds of different compound groups (e.g. carboxylic acids, carbohydrates, amino acids) and a control without added organics. Phytoplankton cultures were added to the plates and incubated for 24 h. Chlorophyll absorbance, chlorophyll fluorescence and number of cells were measured for bogs, Mediterranean Sea, as well as Elbe and Baltic Sea treatments, respectively (tab. S1, tab. S9). Treatments were performed in triplicates. Controls were performed as duplicates (Baltic Sea and bog dataset) or triplicates (Elbe estuary and Mediterranean Sea dataset). The difference in absorbance, fluorescence or cell counts between treatments and control was used as indicative of growth (see detailed explanation in (Martens et al., 2024)). The ecoplates contained 31 dissolved organic compounds, of which 28 were included in our study (tab. S2). D-Xylose, Tween 40, and Tween 80 were not included as their optical properties may interfere with the photometric measurements.

All strains were kept in nutrient enriched media that contained vitamins but no further organic substances and kept in a 12h:12h light:dark cycle with approximately 70 - 150 µmol photons s^-1^ m^-2^ in the light phase (tab. S1). Strains were non-axenic as we know that some of them need their microbiome to survive (Pang et al., 2022; Yao et al., 2019), or reacted sensitively to antibiotics, e.g. *Ostreococcus* (from previous unpublished data) (see also discussion). Detailed information about the laboratory procedures is also given elsewhere (Listmann et al., 2021; Martens et al., 2024).

### Data analysis

We applied a Welch’s t-test (t.test() from stats in R, see below) with p ≤ 0.05 to analyze if measurement values (e.g. cell counts; see above) on the plates were higher in the presence of the different organic compounds than in the control. We consider these significantly positive effects indicative of the use of the respective compounds. Additionally, we analysed if the presence of a compound was associated with at least 10 % higher measurement values than the controls’ mean. This was done to emphasize that many compounds were attributed with elevated measurement values, but were not statistically significant as a result of either small sample sizes (mostly n = 3) or variance. Results are shown in figure 2 and in the supplementary material (fig. S1 - S2, tab. S3 - S5). Moreover, we provide information about the effect size of the significant effects (fig. S3 - S4). Effect size is expressed as % above the controls’ mean. In the bog data, control values were partially negative, which did not allow us to quantify how much higher the treatment values were compared to the control. Here, we only show the qualitative results, i.e., whether treatment values were significantly higher than the control (e.g. fig. 2a).

**Fig. 2:**
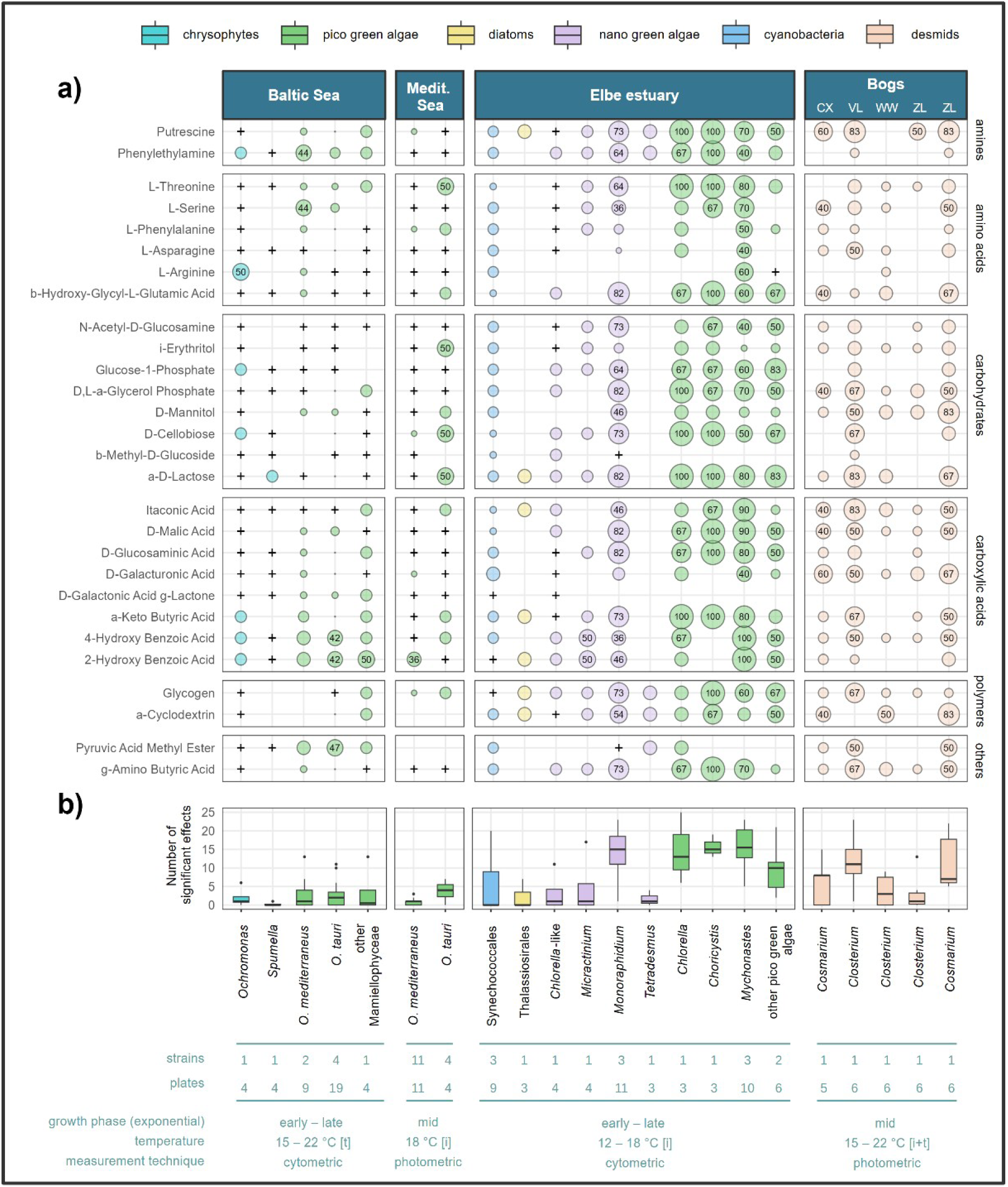
Observed effects in the presence of organic compounds. Points in (a) show the proportion of significant effects (t-test, p ≤ 0.05) across the plates of each phytoplankton group. Numerical values are shown where > 33.3 %. The + signs in (a) indicate measurement values ≥ 10 % higher than the control on at least one plate. This was not obtained for the desmids (see methods). In (b), we show the total number of significant effects per plate. At the bottom, we indicate the number of plates and strains included as well as methodological aspects (fig. 1b). [i] = measurements at the strains’ incubation temperatures, [t] = temperature treatments. CX = Cuxhaven, VL = Vlasina Lake, WW = Wohldorfer Wald and ZL = Zeller Loch. Details see supplementaries (fig. S1 - S6, tab. S1 - S5).

We also applied a Welchs’ t-test (t.test() from stats in R, see below) to analyse differences in the number of significantly positive effects between the different phytoplankton groups in figure 2b (tab. S6).

Of the 28 compounds on the plates, 2 contained P, 11 contained N, and 15 can be understood as sole C sources (tab. S2). For each dataset we calculated the proportion of C, N and P sources that were associated with significant effects across plates.

Data analysis and illustration was carried out in R (version 4.4.1), including the packages stats (version 4.4.1), ggplot2 (3.5.1), svglite (version 2.1.3), patchwork (version 1.3.2.), ggpattern (version 1.1.1), rnaturalearth (version 1.1.0), sf (version 1.0-21), ggspatial (version 1.1.9) and tidyverse (version 2.0.0). We also used LibreOffice Draw (version 7.1.2.2) and google slides for overview figures and the addition of text notes and chatGPT (GPT-5) to streamline R code and to check the finalized manuscript for common grammatical and typographical errors.

## 3. Results

### Observed effects across taxa and habitats

In 41 of 46 strains (89 %), at least one compound on at least one ecoplate was associated with significantly higher measurement values than the control (p ≤ 0.05) (fig. 2a-b, tab. S3 - S5, fig. S1 - S2). On average, 6 of the 28 compounds were associated with significant effects per plate. Five *Ostreococcus* strains from the Mediterranean Sea did not experience significantly increased fluorescence values in the presence of compounds (tab. S4). Notably, all of these had various cases where the fluorescence in the presence of compounds was ≥ 10 % above the control (tab. S4 - S5). Significant effects were found for N and P containing compounds, as well as sole C sources (tab. S5, tab. S2). Effects appeared for every single compound, often across a wide range of taxa and habitats (fig. 2a, tab. S5, fig. S1). For instance, 4-Hydroxy Benzoic Acid was associated with significant effects in 17 of the 22 phytoplankton groups shown in figure 2a, covering all included functional groups and habitats. For some compounds, significant effects rarely appeared across habitats, especially for b-Methyl-D-Glucoside and D-Galactonic Acid g-Lactone (fig. 2a). Also L-Arginine was hardly associated with significant effects (fig. 2a). L-Arginine however was associated with a higher proportion of significant effects across plates in single taxa. This is particularly mentionable for *Mychonastes*, where effects appeared on 60 % of the plates (fig. 2a), and in all three strains (tab. S5, fig. S1).

### Specific results found in the different datasets

In the freshwater habitats, a large proportion of combinations of phytoplankton groups and compounds was associated with significant effects (fig. 2a). The different phytoplankton groups experienced on average around eight significant effects per plate (fig. 2b, tab. S3). In the Elbe estuary dataset, different pico green algae (e.g. *Mychonastes*, *Choricystis*) as well as the nano green alga *Monoraphidium* achieved a significantly higher number of effects than other groups such as *Tetradesmus* and Thalassiosirales (fig. 2b, tab. S6). Moreover, different pico green algae as well as *Monoraphidium* were characterized by more consistent behavior. In other words, when effects appeared, this was often the case on a high number of the included ecoplates, which covered bioreplicates and growth phases. In the case of *Mychonastes* e.g. in up to 100 %, and in *Monoraphidium* in up to 82 % of the plates (fig. 2a). It is particularly mentionable for these taxa, as the plates did also each include three different strains, which are likely different species (tab. S1). Among the bog desmids, *Closterium* from a puddle near Vlasina Lake in Serbia as well as *Cosmarium* from Zeller Loch in Germany behaved more consistently across plates, which included different temperature treatments: Significant effects appeared on up to 83 % of the plates each (fig. 2a). The average number of effects per plate was also higher in these strains compared to the other desmids (fig. 2b, tab. S3), however, due to high variation, differences were only significant between *Closterium* from Serbia and Zeller Loch (tab. S6). The average effect size accompanying significant effects in the Elbe estuary strains was 15 % (fig. S3 - S4, tab. S4 - S5). For the desmids, effect size was not obtained (see methods). In the Elbe estuary, proportions of sole C sources as well as N and P sources associated with significant effects were 18 %, 21 % and 41 %, respectively. Among the bog desmids these were 16 %, 14 % and 22 %.

The number of significant effects observed in the oceanic strains was lower compared to that observed in the freshwater strains, on average two per ecoplate across taxa (fig. 2b, tab. S3). Several phytoplankton groups from the freshwater habitat experienced (e.g. *Mychonastes*, *Closterium* from Serbia) significantly more effects than taxa from the oceanic ecosystems (e.g. *O. tauri* and *O. mediterraneus* from both oceanic habitats) (fig. 2b, tab. S6). For other groups such as Thalassiosirales or the German *Closterium* strains results overlapped with those from the Mediterranean and Baltic Sea taxa (fig. 2b, tab. S6). Where a compound was associated with significant effects in one of the oceanic phytoplankton groups, this was at most the case in 50 % of the included plates (fig. 2a). Variation comes from differential behavior along different growth phases (fig. S5) and temperature treatments (fig. S6) in the Baltic Sea (see also below), as well as different strains of *O. tauri* and *O. mediterraneus* included in each dataset (Mediterranean Sea, Baltic Sea) (fig. S2). *Spumella* achieved a lower number of significant effects than the other Baltic Sea taxa (fig. 2b, tab. S3), but differences between taxa were not significant (tab. S6). Pico green algae from the Baltic Sea - mostly *Ostreococcus* - experienced significant effects particularly often in the presence of different carboxylic acids and less often in the presence of carbohydrates (fig. 2a). Pyruvic Acid Methyl Ester was often associated with significant effects in the Baltic Sea *Ostreococcus* strains, across growth phases and temperature treatments (see also fig. S1), but never across the Mediterranean Sea strains incubated at 18 °C in the mid exponential phase (fig. 2a). Notably, in the Mediterranean Sea, fluorescence also never exceeded the control by ≥ 10 % in the presence of this compound (fig. 2a, fig. S1). In the Mediterranean Sea strains, significant effects appeared less often for *O. tauri* (average 1 per plate) than for *O. mediterraneus* (average 4 per plate) (fig. 2b, tab. S3). However, the difference lacks statistical significance (tab. S6) possibly attributed to the low number of *O. tauri* strains. It is mentionable that many compounds with no significant effects were associated with measurement values exceeding the control by ≥ 10 % in both oceanic habitats (fig. 2a). This was much more often the case than in the Elbe estuary. Generally, effect size was higher in oceanic strains, across the included taxa and functional groups (tab. S5). It was on average 90 % in the Baltic Sea and 76 % in the Mediterranean Sea (fig. S3 - S4, tab. S4 - S5). The proportions of sole C, N, and P sources associated with significant effects were 5 %, 3 % and 0 % in the Mediterranean Sea and 7 %, 6 % and 3 % in the Baltic Sea.

### Role of strains, bioreplicates, growth phase and temperature treatments

As described, the measurements were carried out at different time-points during exponential growth (Elbe estuary, Baltic Sea) and included different temperature treatments (Baltic Sea, bogs). Moreover, some taxonomic groups (*O. tauri*, *O. mediterraneus*, *Mychonastes*, *Monoraphidium, Cosmarium, Closterium*) were included with multiple strains. In some cases, multiple bioreplicates were included (Elbe estuary, bogs). For clarity, results concerning these parameters are shown in the supplementary materials (fig. S1 - S5, tab. S4 - S5, tab. S8). Generally, effects appeared across strains, growth phases, temperatures and replicates (fig. S1 - S2, fig. S5 - S6). In the Baltic Sea, results point towards a negative relationship of significant effects with temperature and growth phase (fig. S5 - S6). Nevertheless, for the other habitats, no clear patterns emerged with respect to these parameters (fig. S5 - S6). Strains from the same taxon often behaved similarly with respect to the average number of significant effects across plates (fig. S2). For instance, all three strains of *Mychonastes* had on average > 10 significant effects (fig. S2, tab. S4, fig. S1). Sometimes differences between strains were more pronounced. For instance, *O. mediterraneus* strain 19.5 from the Baltic Sea hardly experienced significant effects (average < 1) in contrast to strain 4.3 (average 6) (fig. S2, tab. S4, fig. S1). Beyond strain variation, cultures sometimes behaved differently in different bioreplicates. For instance, *Mychonastes* strain P6 achieved 5 to 21 significant effects in the late phase depending on the replicate (fig. S2, tab. S4). Bioreplicate variation was also rather strong in different desmids (fig. S2, tab. S4).

While the methodologies of the datasets vary e.g. in terms of covering different growth phases and temperatures, as well as applying different measurement techniques (see also fig. 1b, tab. S1), there is no indication that these differences significantly affected the observations reported in the present study. In particular, there is no evidence for a mentionable effect on the described differences between different phytoplankton groups. For clarity, this is elaborated in the supplementary material (tab. S8).

## 4. Discussion

### Mixotrophy as a ubiquitous trait in phytoplankton

Mixotrophic behavior in terms of using dissolved organic compounds has been reported for different phytoplankton groups from various locations around the world (fig. 3, tab. S7). Previous studies have provided genetic evidence for phytoplankton possessing the physiological toolkits to use organic compounds (Hagström et al., 2021; Li et al., 2021; Muñoz-Marín et al., 2024; Palenik et al., 2006; Zhang et al., 2021; Zhu et al., 2024). Some studies integrate DOM acquisition into the bigger picture of metabolic processes (Foresi et al., 2024; Villanova & Spetea, 2021). The results of the current synthesis, including growth response data from 46 phytoplankton strains across taxa, functional groups and ecosystems contribute further indication of ubiquitous organic matter acquisition. Unlike proposed in the past, the uptake of organic compounds is not limited to certain essential or limiting substances, such as vitamins, or N and P sources. The observations indicate that mixotrophy is not strictly linked to certain habitats, functional groups or taxa, but rather universal. Previous studies have shown that mixotrophic processes in phytoplankton involve genes commonly found in heterotrophs (e.g. animals, bacteria, fungi), for instance when it comes to membrane transporters such as the urea transporter DUR3 (Li et al., 2021) or the amino acid transporter families SLC7, SLC36, SLC38, and AAPJ (Li et al., 2021; Zhang et al., 2021). Altogether, the universal appearance of mixotrophy and the presence of strictly orthologous genes across domains and lineages imply that heterotrophic processes in phytoplankton represent retained ancestral traits, rather than something that certain phytoplankton taxa acquired over time. Previous studies have shown that many eukaryotic genes are deeply conserved across very distantly related lineages, so that even seemingly unrelated organisms, such as phytoplankton and animals, share homologous genes inherited from their common eukaryotic ancestor (see e.g. (Armbrust et al., 2004; Merchant et al., 2007)). The apparent ubiquity of heterotrophic skills indicates a strong selection pressure imposed on phytoplankton to maintain the ability to use organic compounds (see also (Selosse et al., 2017)).

**Fig. 3:**
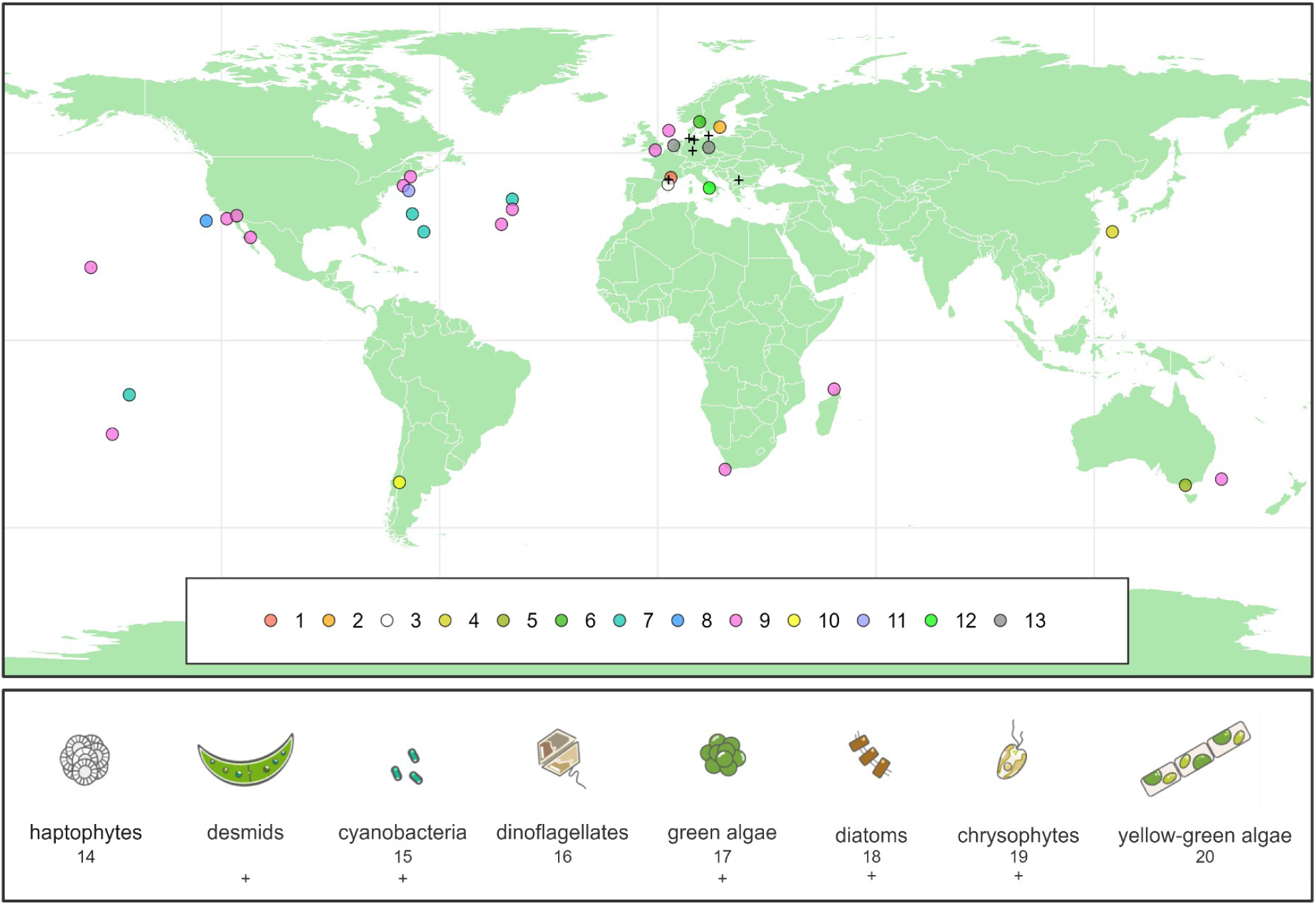
Reported mixotrophic behavior in terms of using dissolved organic matter (non-exhaustive). Data points on the map show the approximate areas from which phytoplankton with reported mixotrophic behavior originated. Below, we show phytoplankton groups with reported mixotrophic behavior. Data from this study (+) and previous research: 1 = (Foresi et al., 2022), 2 = (Hagström et al., 2021), 3 = (Mena et al., 2024), 4 = (Zhang et al., 2021), 5 = (Liu et al., 2024), 6 = (Villanova et al., 2024), 7 = (Muñoz-Marín et al., 2020), 8 = (Palenik et al., 2006), 9 = (Godrijan et al., 2020), 10 = (Beamud et al., 2014), 11 = (Balch et al., 2023), 12 = (Meyer et al., 2022), 13 = (Spijkerman et al., 2017), 14 = (Balch et al., 2023; Beamud et al., 2014; Godrijan et al., 2020), 15 = (Hagström et al., 2021; Muñoz-Marín et al., 2020; Palenik et al., 2006), 16 = (Li et al., 2021; Mena et al., 2024; Zhang et al., 2021), 17 = (Abdullin & Bagmet, 2015; Beamud et al., 2014; Foresi et al., 2022; Hagström et al., 2021; Patnaik & Mallick, 2020; Spijkerman et al., 2017), 18 = (Abdullin & Bagmet, 2015; Mena et al., 2024; Meyer et al., 2022; Villanova et al., 2024; Zhu et al., 2024), 19 = (Hiltunen et al., 2012), 20 = (Liu et al., 2024). See also table S7. A comprehensive map of genes potentially involved in organic matter transport in *Synechococcus* has been provided elsewhere (Muñoz-Marín et al., 2024).

### Acquisition of organic compounds is not strictly linked to specific extrinsic or intrinsic conditions

Mixotrophic behavior has been proposed as a strategy to deal with photosynthetically inconvenient conditions such as nutrient deficiencies or light limitation (Calderini et al., 2022; Foresi et al., 2022; Li et al., 2021; Tuchman et al., 2006). Recent studies show that the expression of membrane transporters for the uptake of organic compounds can be upregulated during N or CO_2_ deficiencies (Li et al., 2021; Zhang et al., 2021). The results of the present study indicate that nutrient and light limitation - conditions not imposed by the experimental design - are not prerequisites for mixotrophy. The finding that sole C sources were used as thoroughly as N and P sources additionally highlights that under the present conditions, N and P acquisition was not the primary factor. A comparably low relevance of the N and P sources can likewise be concluded from former research, which also included phytoplankton strains from rather oligotrophic areas (Godrijan et al., 2020) (see also fig. 3, tab. S7). For some of the Elbe estuary strains, we moreover previously showed that the use of compounds was not increased under light reduction (Martens et al., 2024). Yet, the apparent varying mixotrophic behavior of different strains across growth phases, temperature treatments, and bioreplicates, suggests an influence of intrinsic and extrinsic factors. For instance, as shown in our previous work (Listmann et al., 2021), the more thorough use of compounds by Baltic Sea *Ostreococcus* strains at lower temperatures and in the early exponential phase could be explained by insufficient photosynthetic activity. Nevertheless, in our study, significant effects appeared across growth phases, temperature treatments and bioreplicates. This indicates that specific extrinsic and intrinsic conditions are not a general prerequisite for exploiting heterotrophic pathways. Mixotrophic behavior was particularly consistent in certain taxa such as *Choricystis* and *Mychonastes*, implying constitutive, opportunistic rather than conditional compensatory traits. Phytoplankton are constantly surrounded by DOM in aquatic ecosystems, derived from e.g. terrestrial inputs, exudation and excretion, viral lysis, or sloppy feeding (Jørgensen et al., 2014; Thornton, 2014; Wagner et al., 2020). The use of organic compounds may enhance competitive fitness, explaining its widespread occurrence.

### An attempt at explaining the observed differences across taxa and habitats

The results of the present study imply that different taxa from different habitats have different mixotrophic traits. In other words, available genetic information and the extent to which it is expressed in response to environmental drivers differ. The varying behavior of different taxa per habitat - and similar behavior of strains belonging to the same taxonomic group (e.g. *Mychonastes*, *Ostreococcus*) - indicate that mixotrophic traits are shaped by ancestry. Notably, this was overall more distinct for different genera, and less distinct for different species. The different abilities imply that different phytoplankton groups - in line with the *Plankton Paradox* - occupy different ecological niches. These factors limit interpretations about ecosystem-specific adaptations. There are two taxonomic groups, *Ostreococcus* and *Closterium*, which were isolated from different habitats. While *Ostreococcus* behaved similarly across habitats, and across various strains each, the results from the Serbian *Closterium* strain were particularly distinct from the strains from northern and central Germany. The complying results across *Ostreococcus* from different habitats could indicate a strong ancestral impact, but may also result from similar habitat conditions. Both the Mediterranean Sea and Baltic Sea are semi-enclosed marine habitats, and the sampling areas are rather closely associated with the terrestrial world, resulting in e.g. similar DOM availability in the relevant areas (e.g. western Baltic Sea, Mediterranean lagoons, see also tab. S1) (Amaral et al., 2023; Hoikkala et al., 2015; Lodeiro et al., 2021; Seidel et al., 2017). Notably, one finding particularly distinct between the two oceanic habitats was the apparent exclusive (and rather thorough) use of Pyruvic Acid Methyl Ester among the Baltic Sea *Ostreococcus* strains. The ability to use Pyruvic Acid Methyl Ester - a precursor of Pyruvate - may impose a stronger selection pressure on strains from higher latitudes, in line with lower light availability. However, without further genetic and physiological evidence, this remains speculative. The varying results from *Closterium* isolated from different habitats could indicate habitat-specific adaptations, however, we lack relevant data about the specific bogs and hence cannot draw mechanistic conclusions. As described before, the use of compounds appeared more conditional in some taxa (e.g. *Ostreococcus*), with certain compounds being for instance only used in a certain growth phase per strain - but with high effect sizes. In other taxa, such as *Mychonastes*, use appeared more constitutive, i.e. use across various included ecoplates and growth phases, with lower effect sizes. If we connect these findings to the strains’ origin, use was more conditional in all oceanic strains, and more constitutive in many freshwater strains. Mechanistically, this may be explained by adaptation to the more steady and thorough supply of DOM in bogs and estuaries (Hopple et al., 2019; Novak et al., 2019; Roth et al., 2014; Tobias-Hünefeldt et al., 2024; Watras & Hanson, 2023). Nevertheless, the results should be interpreted with caution, given the small number of taxa included - especially from the oceanic habitats - combined with the fact that different taxa and strains may occupy different ecological niches. Lastly, with respect to data analysis, we should consider that the apparently more conditional and hence variable behavior among the oceanic strains may have affected the number of significant effects observed.

### Potential interactions with bacteria

In our study, strains were kept under non-axenic conditions. Hence, there may be concern about the potential microbial supply with NH_4_ or CO_2_ as a confounding factor (Liu et al., 2024). However, this is not supported by our results. A strong impact of NH_4_ supply would indicate that N sources were used more thoroughly than e.g. sole C sources. A strong impact of microbial CO _2_ would imply effects mostly towards the later exponential phase, where CO_2_ may become rather limiting. Both patterns were not observed. Additionally, in our previous work, we carried out an ampicillin treatment in one of the *Mychonastes* strains (Martens et al., 2024). In figure S7 we provide the associated data analyzed in the same way as the data in the present study. The results with respect to the number of significant effects did not differ between the plates with and without antibiotics (p > 0.05). Given the mentioned points, there is no indication that the microbiome strongly affected our results towards false positive effects. As described before, former studies have provided evidence for the ability of phytoplankton to directly use organic compounds, e.g. based on transcriptomics, compound labeling and growth responses in axenic cultures (Hagström et al., 2021; Li et al., 2021; Liu et al., 2024; Mena et al., 2024; Palenik et al., 2006; Zhang et al., 2021; Zhu et al., 2024). Previous work with axenic cultures has also particularly provided evidence for the microbiome-independent uptake of compounds in different taxa relevant in our study, e.g. *Ostreococcus* (Foresi et al., 2022, 2024), *Mychonastes* (Abdullin & Bagmet, 2015) and *Tetradesmus* (Patnaik & Mallick, 2020). Nevertheless, the microbiome could play a role in various ways, e.g. in the breakdown of larger compounds, such as the polymers. Also, as described in our previous work, there is the possibility that some positive effects of organic compounds on phytoplankton growth are masked by negative effects imposed on phytoplankton by e.g. microbial toxin production (Martens et al., 2024).

## 5. Conclusion

Understanding phytoplankton’s metabolic processes and environmental drivers is fundamental to determine their functions in ecosystems and properly map their role in climate change. The present synthesis highlights the prevalence of mixotrophy in terms of dissolved organic compounds in phytoplankton across taxa and habitats. Our results suggest that future experimental design in phytoplankton research must move beyond the strict autotrophic paradigm to account for carbon flux from dissolved organic sources. With DOM being one of the largest carbon pools on Earth (Hansell et al., 2009; Houghton, 2003), and phytoplankton being key mediators of carbon sequestration (Basu & Mackey, 2018), this study underscores the ecological significance of heterogeneous nutritional strategies and the necessity to address existing knowledge gaps to finally draw conclusions about the impact of mixotrophy in the context of single ecosystems, and on global scales.

## Acknowledgements

We thank Klaus von Schwartzenberg, Sigrid Körner, and Gwenael Piganeau for providing different phytoplankton strains, Franziska Kerl and Emilia Ehlert for their contributions to the laboratory experiments, and Stefanie Schnell for general laboratory support.

## Funding

This work was supported by the Deutsche Forschungsgemeinschaft (DFG, German Research Foundation) [grant number: 407270017 (RTG2530)]. We acknowledge financial support from the Open Access Publication Fund of Universität Hamburg.

## Conflict of Interest

We declare that we have no competing interests.

## Data availability statement

The supplementary materials show the raw measurement values (e.g. cell counts; tab. S9), all numeric results referred to in the manuscript (e.g. p-values, effect sizes; tab. S3 - S6) as well as metadata (tab. S1). Further raw data and metadata have been published elsewhere for the Elbe estuary (https://doi.org/10.25592/uhhfdm.18666), the Baltic Sea (https://zenodo.org/records/3958784), and the Mediterranean Sea (https://doi.org/10.25592/uhhfdm.18670).

## Material from previous publications

We hold the copyright for all included datasets, including those previously published.

## Notes

### Competing Interest Statement

The authors have declared no competing interest.

### Summary of Updates

The manuscript was completely revised including statistical methods and interpretations.

