## Supplementary figures for "Mixotrophy in phytoplankton: Prevalent use of organic compounds and lineage-specific variations suggest conserved ancestral traits"

**Fig. S1: Significant effects (green; t-test, p ≤ 0.05) of each compound on each ecoplate. The + indicates where the mean measurement value of the 3 well replicates was ≥ 10 % higher than the control.**

The horizontal scales shows the different compounds (bottom) sorted by compound group (top). On the vertical axis we show the different plates which include different taxa, strains, incubation times, temperatures, growth phases and replicates. Precisely, it is named as such: >>taxon – strain – temperature – growth phase (1 – 3 = early - late) – replicate<<. The vertical panels show different ecosystems (BS = Baltic Sea, MS = Mediterranean Sea, EE = Elbe estuary, Bo = Bogs) and functional groups (Des = desmids, Chry = chrysophytes, PG = pico green algae, NG = nano green algae, Dia = diatoms, Cya = cyanobacteria) (see also tab. S1). For the bog desmids, different origin is indicated (ZL = Zeller Loch, VL = Puddly near Vlasina Lake, WW = Wohldorfer Wald, CX = Cuxhaven). Note that effect size was not obtained for the bog dataset, explaining the absence of + here.

green = significant effect

**white** = no significant effect

**+** = mean measurement value of the 3 well replicates ≥ 10 % higher than the control

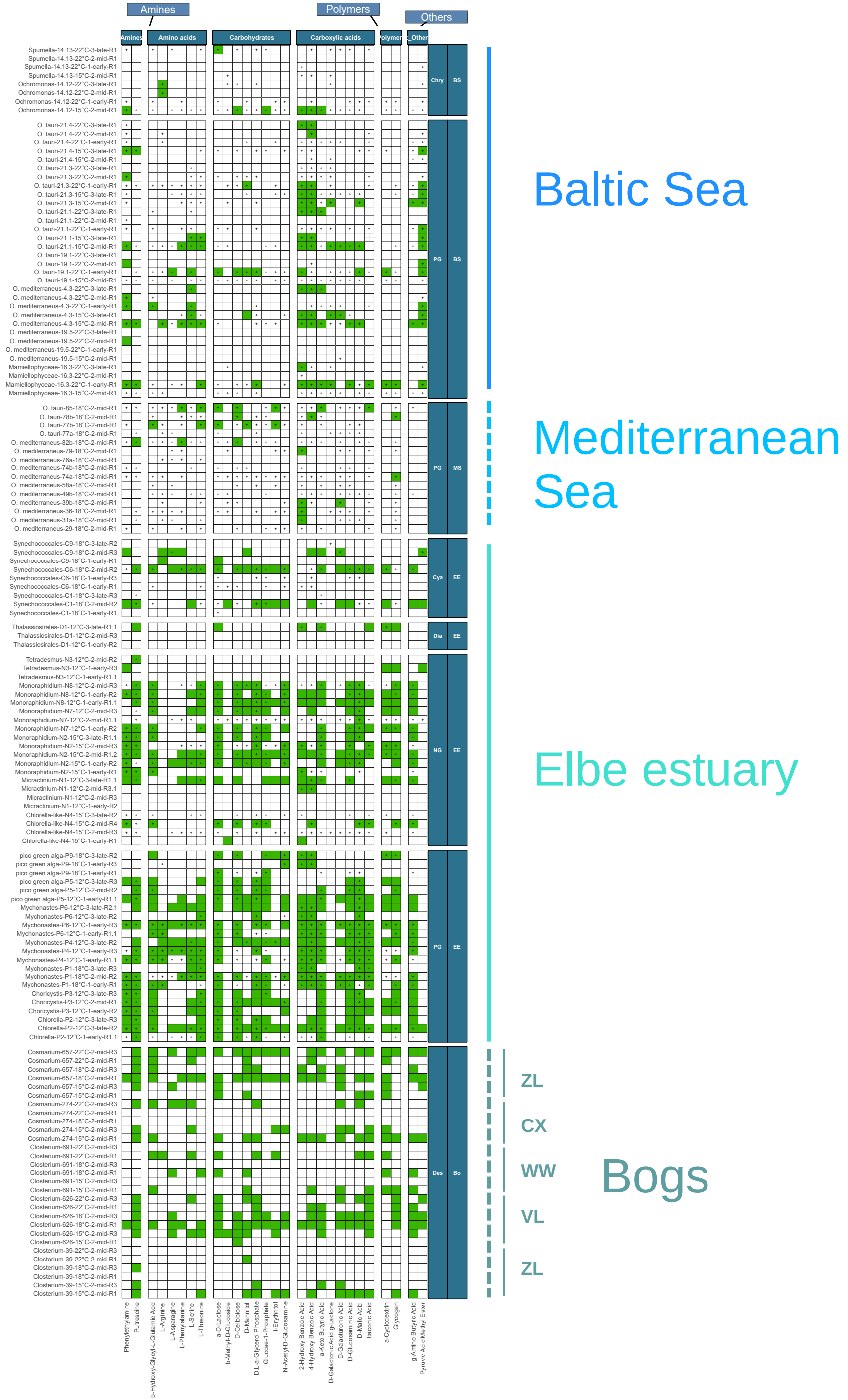

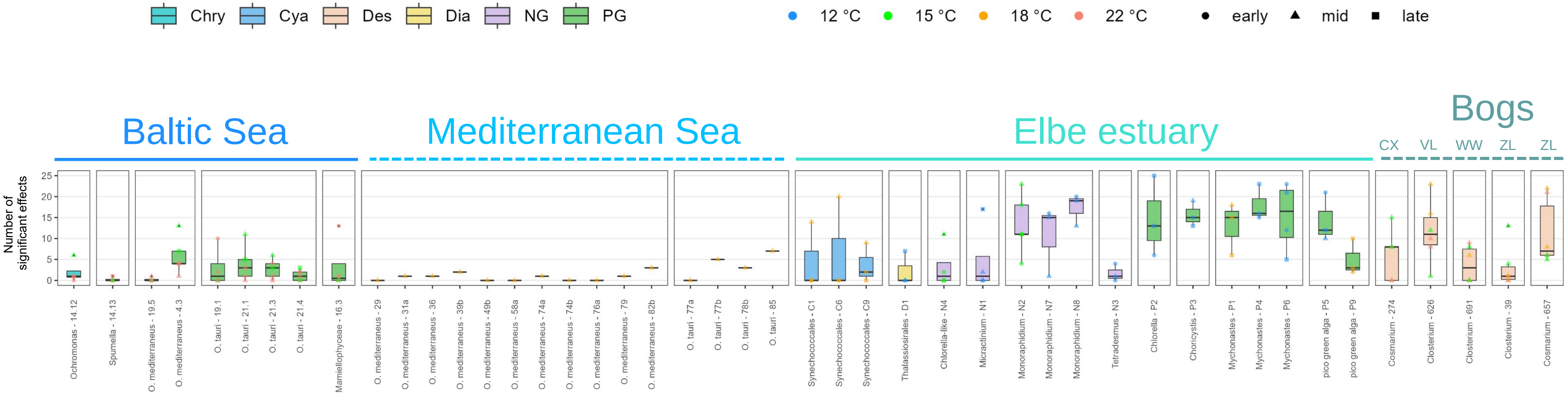


**Fig. S2: Number of significant effects (t-test, p ≤ 0.05) per ecoplate clustered by strains covering different taxa from different habitats.** Color scheme of the boxplots indicates functional group (Des = desmids, Chry = chrysophytes, PG = pico green algae, NG = nano green algae, Dia = diatoms, Cya = cyanobacteria). Color scheme of the points indicate the experimental temperature. Point shape shows the timepoint during the exponential phase. Note that effect size was not obtained for the bog dataset, explaining the absence of ‘+’ here. For the bog desmids, different origin is indicated (ZL = Zeller Loch, VL = Puddly near Vlasina Lake, WW = Wohldorfer Wald, CX = Cuxhaven).

**Fig. S3: Effect sizes of each compounds with significant effects (t-test, p ≤ 0.05) per ecoplate.**

Effect size shows how much higher the measurement value (cell count, fluorescence) was in the presence of organic compounds compared to the control [%]. It is shown as the mean across the three well replicates each. The horizontal scales shows the different compounds (bottom) sorted by compound group (top). On the vertical axis we show the different plates which include different taxa, strains, incubation times, temperatures, growth phases and replicates. Precisely, it is named as such: >>taxon – strain – temperature – growth phase (1 – 3 = early - late) – replicate<<. The vertical panels show different ecosystems (BS = Baltic Sea, MS = Mediterranean Sea, EE = Elbe estuary) and functional groups (Chry = chrysophytes, PG = pico green algae, NG = nano green algae, Dia = diatoms, Cya = cyanobacteria) (see also tab. S1). Effect size was not obtained for the bog dataset.

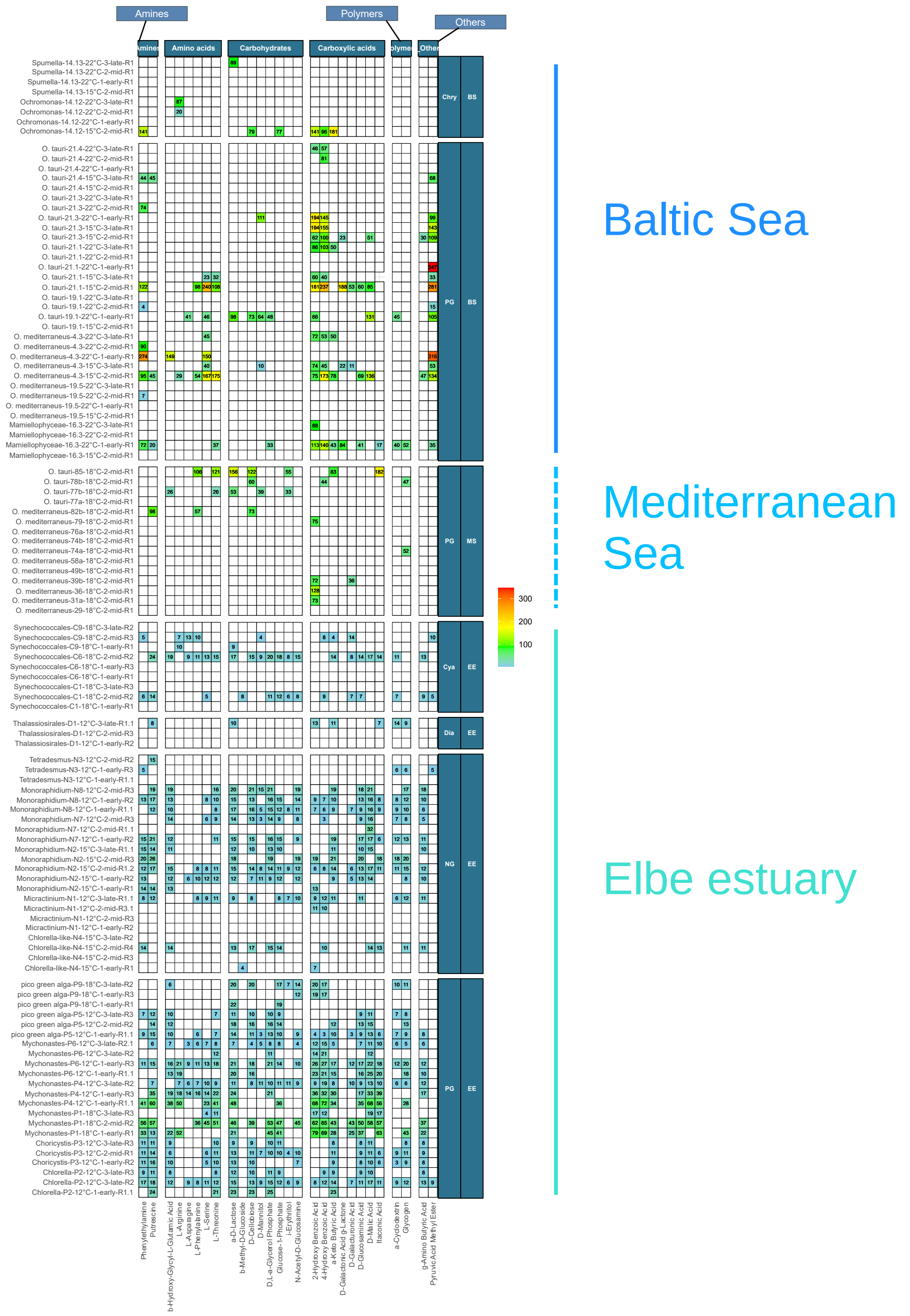

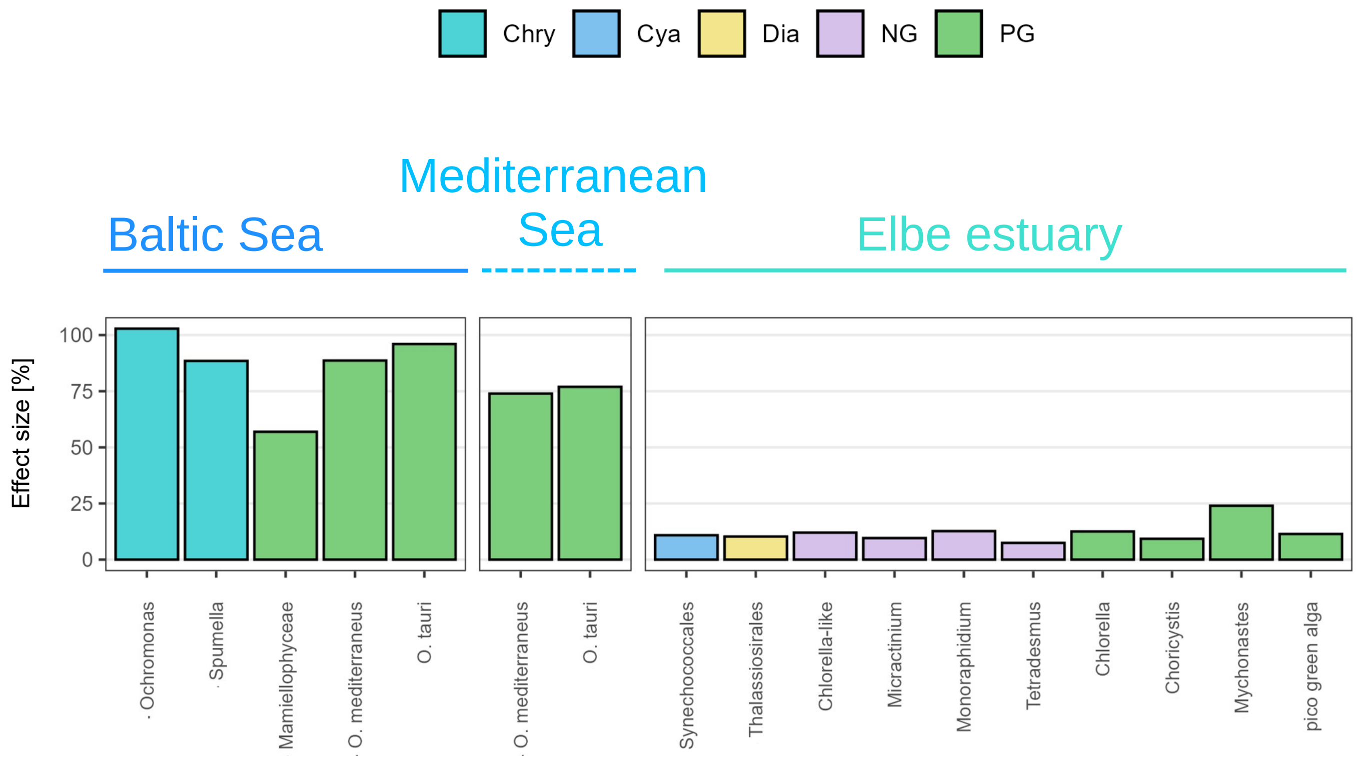


**Fig. S4: Average effect sizes of significant effects (t-test, p ≤ 0.05) per phytoplankton group.** Effect size shows how much higher the measurement value (cell count, fluorescence) was in the presence of organic compounds compared to the control. Color scheme indicates functional group (Chry = chrysophytes, PG = pico green algae, NG = nano green algae, Dia = diatoms, Cya = cyanobacteria). Note that effect size was not obtained for the bog dataset.

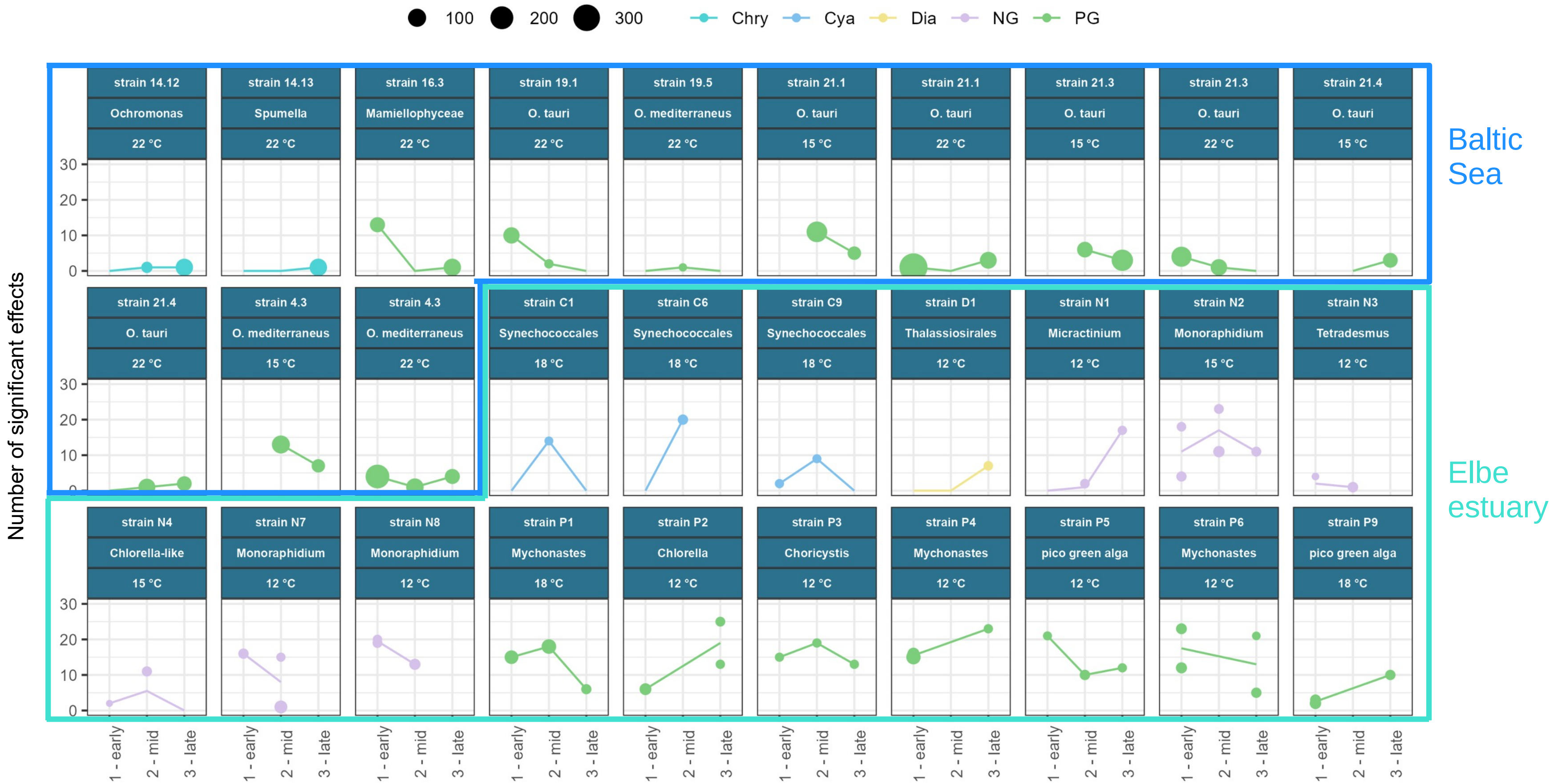


**Fig. S5: Number of significant effects (t-test, p ≤ 0.05) along growth phases (1 = early, 2 = mid, 3 = late exponential).** Panels show the different taxa, strains (e.g. 16.3) and temperatures. Color scheme indicates functional group (Chry = chrysophytes, PG = pico green algae, NG = nano green algae, Dia = diatoms, Cya = cyanobacteria). Data points are connected by a simple line as a visual aid. Point size shows the mean effect size of significant effects per plate [%] as indicated above (no significant effects = no point). Where more than one bioreplicate was included, line is drawn through the mean. Strain and temperature combinations that did not cover multiple growth phases were removed, which also concerns the complete bog and Mediterranean Sea data.

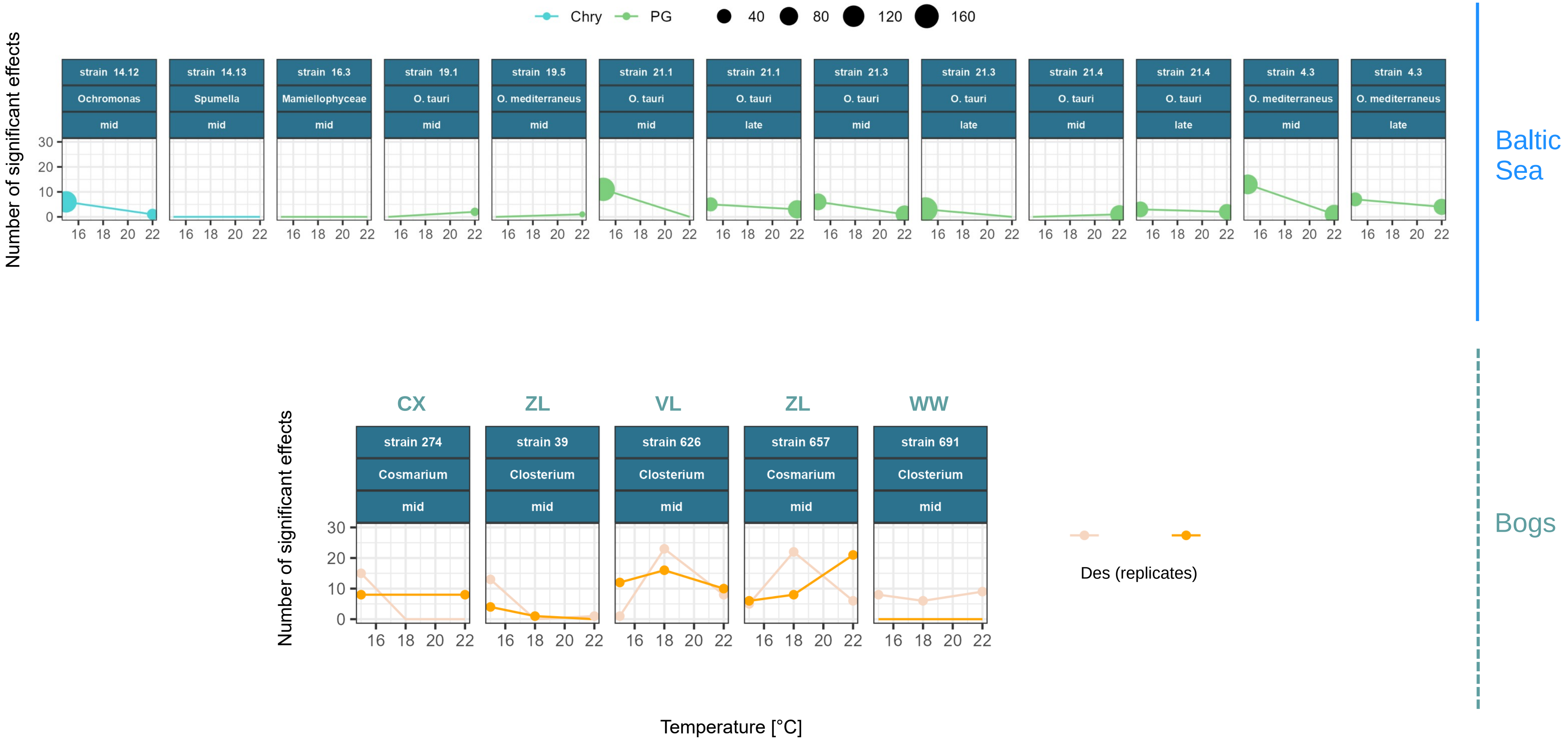


**Fig. S6: Number of significant effects (t-test, p ≤ 0.05) along temperatures.** Panels show the different taxa, strains (e.g. 14.12) and growth phases. Colors indicate the functional group (Chry = chrysophytes, PG = pico green algae, Des = desmids), and for the desmids also different bioreplicates. Data points are connected by a simple line as a visual aid. For the Baltic Sea data point size shows the mean effect size of significant effects per plate [%] as indicated above. For the bog data point size is fixed where effects appeared, as effec size was not obtained. In both datasets, no point indicates no significant effects. Strain and growth phase combinations that did not cover multiple temperatures were removed, which also concerns the complete Elbe estuary and Mediterranean Sea data. For the bog desmids, different origin is indicated (ZL = Zeller Loch, VL = Puddly near Vlasina Lake, WW = Wohldorfer Wald, CX = Cuxhaven).

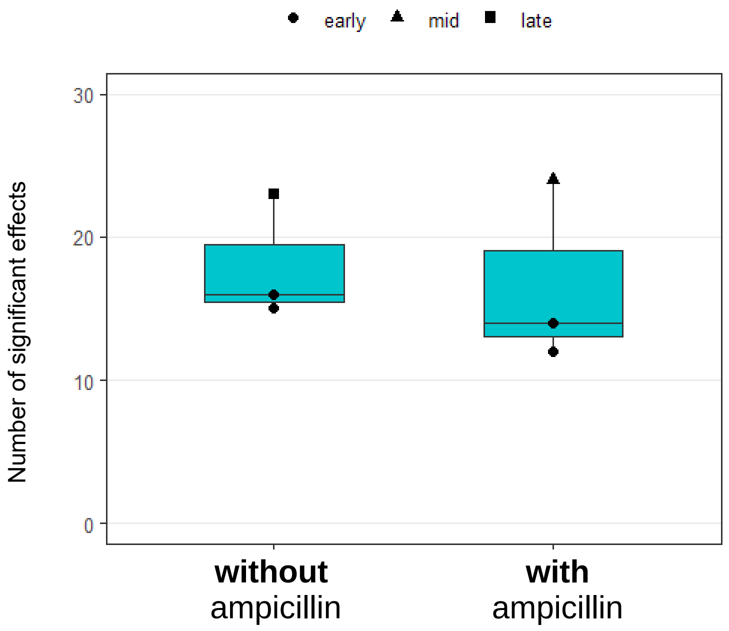


**Fig. S7: Number of significant effects (t-test, p ≤ 0.05) in *Mychonastes* strain P4 from the Elbe estuary with and without addition of ampicillin.** Points show the different included plates which cover different growth phases, indicated by shape. Differences between the two treatments are not significant (t-test, p > 0.05). Mean values are 18 (without ampicillin) and 17 (with ampicilllin). Details see publication 10.1098/rspb.2023.2713.
